# Cytosolic peroxiredoxin Tsa1 influences acetic acid metabolism and pH homeostasis in wine yeasts

**DOI:** 10.1101/2024.12.23.629978

**Authors:** Víctor Garrigós, Cecilia Picazo, Lisa Dengler, Jennifer C. Ewald, Emilia Matallana, Agustín Aranda

## Abstract

Acetic acid is a key metabolite in yeast fermentation, influencing wine quality through its role in volatile acidity. In *Saccharomyces cerevisiae*, acetic acid production involves aldehyde dehydrogenases, primarily Ald6 during fermentation and Ald4 under respiration. However, the regulatory mechanisms of these enzymes throughout fermentation and how they differ in commonly used strains remain partially unclear. This study explores the cytosolic peroxiredoxin Tsa1 as a novel regulator of acetic acid metabolism. *TSA1* gene deletion revealed strain-dependent effects on acetic acid metabolism and tolerance, showing reduced production and enhanced consumption in laboratory media. Under respiration, Ald4-driven acetic acid production, which raises extracellular pH, was mitigated by the absence of Tsa1. During wine fermentation, *TSA1* deletion decreased the initial acetic acid surge by downregulating *ALD6* transcription and enzymatic activity. These findings position Tsa1 as a metabolic regulator and a potential target for modulating acetic acid levels to manage volatile acidity and wine quality.

## Introduction

Volatile acidity is a relevant parameter of wine that derives from acetic acid and related compounds, either in their free form or bound as salts, as outlined in the OIV resolution OIV-OENO 662C-2022. Its main component is acetic acid, which causes a pungent smell associated with vinegar and wine spoilage (Ribéreau-Gayon et al., 2006). The yeast *Saccharomyces cerevisiae* can produce its own share of acetic acid during regular grape juice fermentation (Vilela-Moura et al., 2011). Under respiratory conditions, acetyl-CoA is produced directly from pyruvate by the mitochondrial pyruvate dehydrogenase (PDH) complex. This complex is inactive during grape juice fermentation, so the cytosolic acetyl-CoA required for processes such as lipid synthesis is produced via the PDH bypass pathway. First, pyruvate from glycolysis is decarboxylated to acetaldehyde by pyruvate decarboxylase (PDC). While most acetaldehyde is reduced to ethanol by alcohol dehydrogenases (ADHs) to maintain redox balance, a fraction is further oxidized to acetic acid by aldehyde dehydrogenases (ALDHs). Acetic acid is subsequently converted into acetyl-CoA through the action of acetyl-CoA synthases (ACS), a reaction that requires ATP. There are a variety of ALDH activities described in yeast, distributed in two cellular locations (Navarro-Aviño et al., 1999). Cytosolic ALDHs include Ald6 (the major one), and the stress-induced Ald2 and Ald3, while mitochondrial ALDHs are encoded by *ALD4* (the major one) and *ALD5*. Ald6 is dependent on magnesium ions, while Ald4 uses potassium, and *ALD4* gene is glucose repressed while *ALD6* is fully active during fermentation (Aranda and Ml, 2003). That would explain that during grape juice fermentation, the role of cytosolic isozyme Ald6 in the production of acetic acid is more relevant than its mitochondrial counterpart Ald4 (Remize et al., 2000).

Acetic acid production in *S. cerevisiae* depends on many environmental factors, including the nitrogen and vitamin composition of the grape juice, the initial sugar concentration, and physical parameters like temperature (Vilela-Moura et al., 2011). Oxygen plays a pivotal role in its production too. While aeration has been a tool to reduce ethanol content, it can activate respiratory pathways, leading to an undesirable increase in volatile acidity (Tronchoni et al., 2022). Ethanol reduction is a relevant challenge in modern enology, as rising sugar levels in grapes due to global warming result in elevated alcohol levels in wine (Orduna, 2010). Efforts to divert glycolytic flux toward glycerol production by genetic manipulation have shown potential for reducing these ethanol levels. However, these modifications also lead redox unbalances, leading to an unpleasant rise in acetic acid production that requires additional deletion of ALDH genes (Zhao et al., 2015). Therefore, the regulatory mechanisms governing acetic acid metabolism are of great interest to winemaking researchers to tackle the dual challenges of managing alcohol content and volatile acidity.

Previous works from our laboratory have demonstrated that redox signaling influences metabolic processes under winemaking conditions (Garrigós et al., 2020; Picazo et al., 2019, 2018). One of the main redox systems responsible for oxidative damage repair and cellular redox control is the cytosolic peroxiredoxin-thioredoxin-thioredoxin reductase system (Morano et al., 2012). At the core of this system is Tsa1, the principal cytosolic peroxiredoxin, which detoxifies a range of peroxides, including hydrogen peroxide (H_2_O_2_). The catalytic mechanism of Tsa1 involves the formation of a disulfide bond between its peroxidatic cysteine (C_P_) and resolving cysteine (C_R_) during peroxide reduction. The disulfide bond is subsequently reduced by the cytosolic thioredoxins Trx1/2, which are regenerated by thioredoxin reductase (Trr1) using NADPH as an electron donor. Beyond its role in peroxide detoxification, Tsa1 also acts as a chaperone (Jang et al., 2004) and as a regulator of the cAMP-protein kinase A (PKA) signaling pathway through direct redox control of its catalytic subunits, linking redox regulation to cellular metabolism and aging (Roger et al., 2020). Hence, this redox system plays a crucial role in repairing oxidative damage in key cysteines across a variety of enzymes (Toledano and Huang, 2016), further highlighting its potential involvement in metabolic regulation. For instance, *TRR1* deletion affects the TCA cycle and increases the content of proteogenic amino acids (Picazo et al., 2018). The *trx1*Δ*trx*2Δ double mutant exhibits diminished glycolytic activity but increased lipid synthesis and amino acid metabolism during wine fermentation (Picazo et al., 2019). Additionally, *TSA1* deletion decreases acetic acid production during grape juice fermentation and enhances trehalose accumulation at the end of biomass propagation in molasses (Garrigós et al., 2020).

This manuscript provides a detailed examination of the role of Tsa1 in acetic acid metabolism. *TSA1* and *TRR1* deletion alters acetic acid production and consumption in a manner that varies across different growth conditions and media. In addition, genetic diversity among commercial wine strains contributes to differential tolerance to acetic acid. Cytosolic magnesium-dependent ALDH activity is controlled by Tsa1 during the initial hours of grape juice fermentation. However, mitochondrial potassium-dependent ALDH activity is also influenced by the absence of the *TSA1* gene under low-glucose, respiratory conditions. Therefore, these findings highlight the complexity of acetic acid metabolism regulation through the redox-controlling peroxiredoxin-thioredoxin system.

## Materials and methods

### Yeast strains, growth media and conditions

The yeast strains herein used are listed in Supplementary Table 1. Deletion mutants were generated in the laboratory strain W303 and the haploid wine strain C9 (Walker et al., 2003) using the reusable kanMX marker. This marker was amplified via PCR from the pUG6 plasmid (Guldener et al., 1996) and contains loxP sequences for excision by the Cre recombinase from the YEp-cre-cyh plasmid (Delneri et al., 2000). In the industrial wine strains L2056, T73, and EC1118, CRISPR-Cas9 deletions of the *TSA1* and *TRR1* genes were performed by CRISPR-Cas9 using the pRCC-K plasmid, a gift from Eckhard Boles (Addgene plasmid #81191), and following the protocol described (Generoso et al., 2016). Yeast transformations were carried out using the lithium acetate method (Gietz and Woods, 2002).

For maintenance and standard propagation purposes, yeast cultures were grown at 30 °C in rich YPD medium (1% yeast extract, 2% bactopeptone, 2% glucose). Solid media were prepared by supplementing with 2% agar, and geneticin at a concentration of 200 mg/L was used for selecting kanMX transformants. To evaluate the phenotypic traits of the strains under alternative standard laboratory conditions, cells were cultivated in Glucose Minimal Media (GMM) as previously described (Dengler et al., 2021). Minimal medium SD contained 0.17% yeast nitrogen base, 0.5% ammonium sulfate and 2% glucose.

For growth spot analysis, serial dilutions were prepared from stationary-phase cultures grown in YPD, and 5 μL drops were placed onto selective media. When required, the carbon source in YPD was substituted with 2% fructose, sucrose, glycerol, or ammonium acetate. For stress analysis, SD medium was used adding 0.17% acetic acid or 10% ethanol.

To investigate the impact of yeast colony growth on extracellular pH, solid growth medium was prepared as described (Baron et al., 2013). Extracellular pH was monitored using the “giant colony” growth method (Palkova et al., 1997). An overnight culture grown in YPD was diluted to an OD (600 nm) of 0.01, and 10 µL were spotted onto plates and incubated at 30 °C. To quantitatively investigate the impact of metabolic transitions on extracellular pH, yeast cells were cultured under low-glucose conditions. Cultures were inoculated at an initial OD (600 nm) of 0.05 and incubated at 30 °C with shaking at 220 rpm for 24 hours. The liquid medium contained 1% yeast extract and 0.2% glucose, with the pH adjusted to 5.5 using HCl. Extracellular pH was determined using a colorimetric assay employing the pH-sensitive dye BCP (Baron et al., 2013).

Bioreactor-scale assays were conducted using Synthetic Molasses (SM), prepared as previously described (Garrigós et al., 2020), in a 5LJL ez2-Control bioreactor (Applikon Biotechnology, Netherlands) equipped with proportional, integral and derivative (PID) control units for pH, temperature, oxygen and agitation speed. The bioreactor containing 3LJL of sterilized SM was inoculated with an initial OD (600 nm) of 0.1 from YPD precultures.

Wine fermentation experiments were performed using synthetic must (MS300) prepared as described (Riou et al., 1997). Cells from 2-day YPD cultures were inoculated into 30 mL of must in conical centrifuge tubes at a final concentration of 10^6 cells/mL and the cultures were incubated at 25 °C with low agitation (50 rpm). The vinification progress was monitored by measuring sugar consumption using the dinitro-3,5-salicylic acid (DNS) according to Miller’s method (Robyt and Whelan, 1972).

### Metabolite measurements

Acetic acid and ammonia concentrations were determined spectrophotometrically using coupled enzymatic reactions linked to the NAD⁺/NADH and NADP⁺/NADPH redox pairs. Commercial kits from Megazyme Ltd (Bray, Ireland, www.megazyme.com; K-ACET and K-AMIAR kits) were employed for these measurements, following the manufacturer’s protocols. HPLC was employed to quantify acetic acid in the GMM samples. HPLC analysis was performed on a Shimadzu HPLC system equipped with a Repromer H column from Dr. Maisch GmbH, Germany. A sample volume of 10 µL was injected onto the column using an autosampler cooled to 4 °C. Metabolites were eluted isocratically with 5 mM H2SO4 (Sulfuric Acid Solution, 49 to 51%, For HPLC, Honeywell Fluka). Samples were run with a flow rate of 1.0 ml/min for 25 min at 40 °C. The analytes were monitored using a refractive index detector (Shimadzu RID-20A).

### Measurement of Enzymatic activities

Cytosolic and mitochondrial aldehyde dehydrogenase activities were measured using cell extracts from cultures grown under the winemaking conditions described above. Cell extracts were prepared by washing and resuspending yeast cells in 0.5 mL of a buffer containing 10 mM sodium phosphate (pH 7.0), 1 mM dithiothreitol, and 5% (w/v) glycerol. Cell disruption was achieved with glass beads using a Precellys® Evolution homogenizer, operated for three 20-second intervals at a speed of 4.5 m/s, with 1-minute cooling periods between each interval. Protein concentration was measured using the DC Protein Assay (BioRad), following the manufacturer’s protocol. Cytosolic and mitochondrial aldehyde dehydrogenase activities were assessed (Aranda and Ml, 2003). In both cases, the reaction was initiated by the addition of acetaldehyde, and the increase in absorbance at 340 nm was monitored using a Varioskan LUX microplate reader (Thermo Fisher Scientific).

### Relative gene expression level quantification by real-time PCR and Western Blot

To quantify the relative expression levels of the target genes, yeast cells were harvested at the specified time points, and total RNA was extracted as described previously (Carrasco et al., 2001). The extracted RNA was reverse transcribed using the NZY First-Strand cDNA Synthesis Kit (Nzytech). Gene-specific primers (Zhang et al., 2022) were used to amplify *ALD4* and A*LD6*. The housekeeping gene *ACT1* served as an internal control. Quantitative PCR (qPCR) was performed using the NZYSpeedy qPCR Green Master Mix (Nzytech) on a QuantStudio 3 instrument, following the manufacturer’s protocol. Each reaction was conducted in triplicate, and the average cycle threshold (Ct) value from triplicate samples was used for analysis. Relative transcript levels were calculated using the 2⁻ΔΔCt method. For Western Blot, cells were lysed using glass beads, and whole-cell extracts were prepared in lysis buffer containing 1 M Tris-HCl (pH 7.5), 5 M NaCl, 1 M MgCl₂, 10% (v/v) NP40, 0.1 M PMSF, and a commercial protease inhibitor tablet (complete Mini, EDTA-free; Roche). Proteins were resolved in an SDS-PAGE gel using an Invitrogen mini-gel system and transferred to PVDF membranes using a Novex semi-dry transfer system (Invitrogen, Carlsbad, CA, USA). Immunodetection was performed with the anti-2-Cys-Prx (Abcam) or anti-Prx-SO3 (Abcam) antibodies. Anti-Pgk1 (Invitrogen) was used as the loading control. Signal detection was carried out using the ECL Western Blotting Detection System (GE Healthcare) following the manufacturer’s instructions.

## Results

### *TSA1* deletion has an impact in metabolism of wine yeasts

Previous results indicated that deletion of peroxiredoxin *TSA1* and heterozygous deletion of one of the copies of thioredoxin reductase *TRR1* reduced the amount of acetic acid during winemaking (Garrigós et al., 2020). To determine if this mutation influences acetic acid production in a completely different and industrially relevant environment, the same mutants were tested during growth in synthetic molasses in a bioreactor (Figure 1). This media mimics the composition of industrial molasses used for yeast biomass propagation, albeit with a reduced sucrose concentration to facilitate early investigation of the transition to respiratory metabolism. As previously described, under these conditions, although the *tsa1*Δ mutant cells reached saturation after 24h of growth, it was unable to achieve the same cell density as the parental strain and remained so even during the fed-batch phase. The *trr1*Δ/*TRR1* mutant showed a similar growth profile to the wild-type strain (Garrigós et al., 2020). During the growth in *batch*, at 6 hours, the amount of acetic acid is low and slightly reduced in the mutants. Acetic acid accumulates to reach a maximum at 24 hours in all strains. At this point, the acetate levels in both mutants are comparable to those observed in the wild-type strain. After 24 hours, sucrose is fully consumed (data not shown), and acetic acid decreases in the wild-type strain, reaching a minimum at 51 hours. In contrast, acetic acid levels in the *tsa1*Δ mutant remain elevated during this period. In the *trr1*Δ/*TRR1* mutant, acetic acid reduction is delayed and incomplete compared to the wild type. After 51 hours, fed-batch growth was initiated to maintain yeast’s respiratory metabolism. During this final stage of biomass production, acetic acid levels increase in both the wild type and the *trr1*Δ*/TRR1* mutant, although it starts earlier in the mentioned mutant. Notably, the levels in the *tsa1*Δ mutant remain high throughout the process. These findings indicate that Tsa1 has a relevant impact on acetic acid metabolism during sucrose fermentation, with an effect that appears to differ from its role during microvinification, potentially acting in the opposite direction.

**Figure 1.**
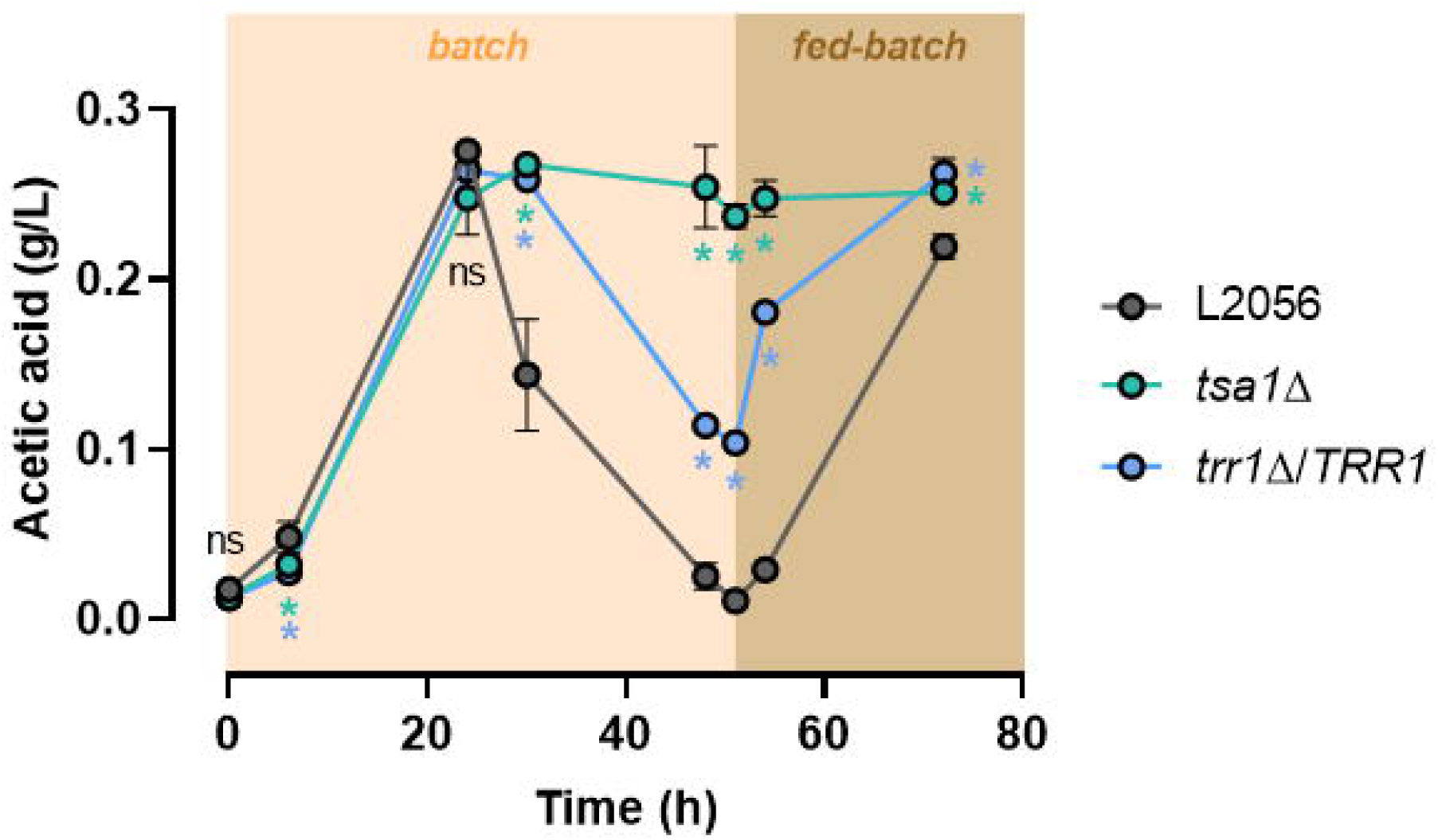
*TSA1* deletion increases acetic acid levels during molasses fermentation. L2056, L2056 *tsa1*Δ and L2056 *trr1*Δ*/TRR1* strains were grown in Synthetic Molasses Medium with 2% sucrose in a bioreactor in batch (up to 51 hours) and fed-batch conditions. Acetic was measured by triplicate and the mean and standard deviation are shown. Significant differences (*p < 0.05, Student’s t-test) between the mutants and the wild-type strain are shown.

As volatile acidity is a relevant phenotype during winemaking and its production varies among commercial wine strains, our next goal was to determine the comparative impact of *TSA1* and *TRR1* mutations on carbon metabolism and stress tolerance across several commercial wine yeasts. To this end, we developed a CRISPR-Cas9 method capable of efficiently deleting all copies of *TSA1* and *TRR1* simultaneously, addressing the diploid nature of most commercial strains. In addition to L2056, we included EC1118, a widely used commercial strain, and T73, another reference strain used in our laboratory.

To test the growth of these strains and their corresponding *TSA1* and *TRR1* mutants, spot assays were performed in different substrates (Figure 2A). In the standard rich glucose-containing medium YPD, *TSA1* deletion does not impact growth. As expected (Picazo et al., 2018), *TRR1* deletion did, particularly in the L2056 genetic background. That may be the reason why the homozygous mutant was not obtained in L2056 strain using the classical homologous recombination approach, and an heterozygotic mutant was used (Garrigós et al., 2020). Similar results were observed when using other easy-to-assimilate hexoses like fructose (YPF) and a disaccharide like sucrose (YPS), the latter commonly used in yeast biomass propagation. Therefore, *TSA1* deletion does not impact growth on fermentable sugars. When a non-fermentable, respiratory carbon source such as glycerol was tested, a similar pattern was observed: *tsa1*Δ mutants displayed no additional phenotype changes. This suggests that *TSA1* deletion does not broadly affect either fermentable or respiratory metabolism.

**Figure 2.**
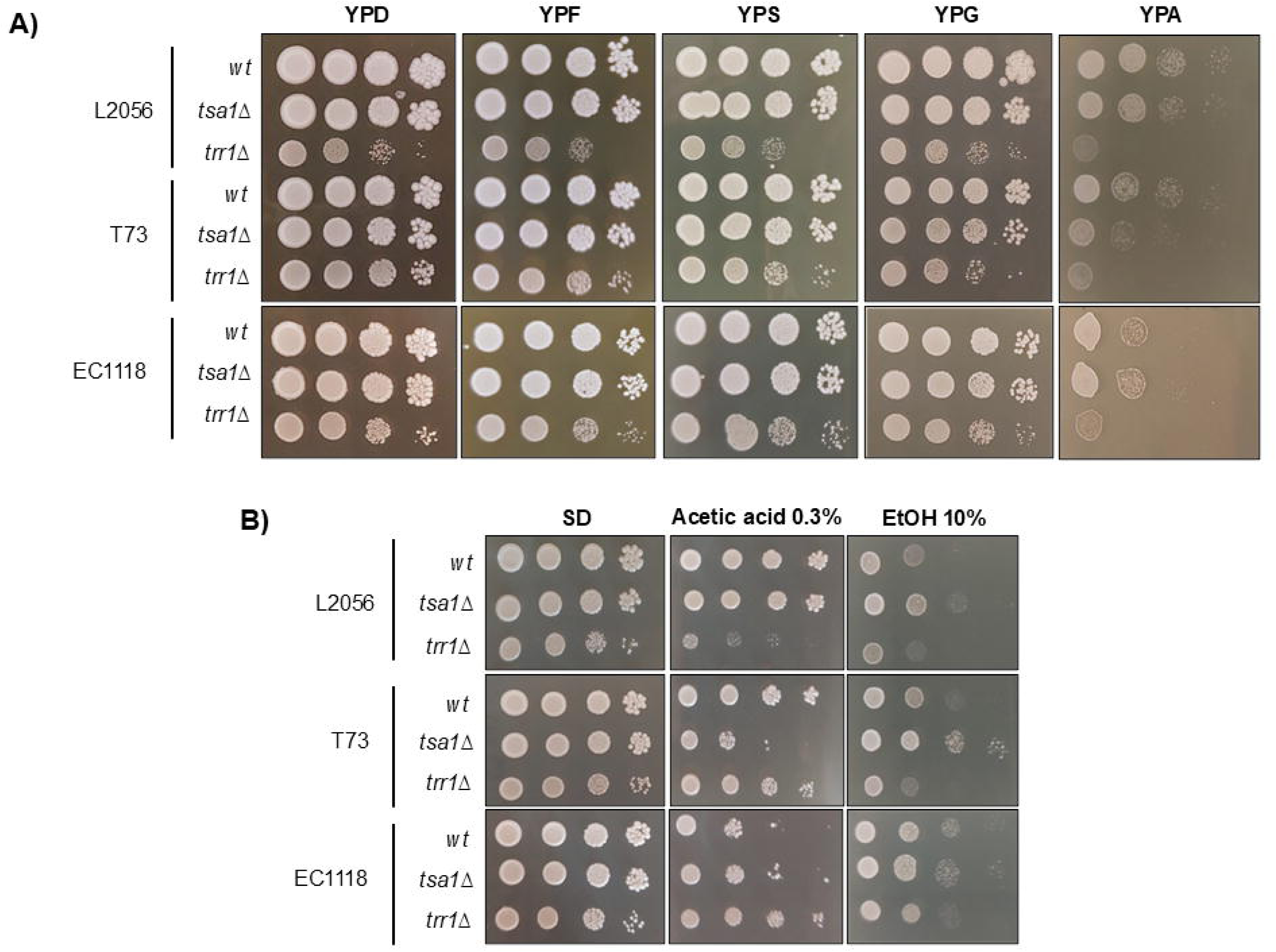
Mutations in *TSA1* and *TRR1* show phenotypic dependance on genetic background. CRISPR-Cas9 deletions of *TSA1* and *TRR1* genes in the commercial wine strains L2056, T73 and EC1118 were tested in different conditions by drop assay. Stationary cultures in YPD were normalized according to OD_600_, serial dilutions were performed, and 5 ml drops were placed in different media. A) Growth in plates of rich media with different carbon sources: 2% glucose (YPD), fructose (YPF), sucrose (YPS), glycerol (YPG) and ammonium acetate (YPA) were used. B) Stress caused by 0.3% acetic acid and 10% ethanol in minimal medium SD plates were performed.

However, when acetate was used as the sole carbon source, growth was generally delayed, particularly for *TRR1* deletion mutants. Interestingly, while *tsa1*Δ mutants in L2056 and EC1118 strains behaved similarly to their parental strains, T73 *tsa1*Δ showed delayed growth, indicating that *TSA1* plays a more significant role in acetate assimilation in this genetic background. Next, the impact of a stressful amount of acetic acid was tested (Figure 2B). This experiment is usually conducted in glucose-containing minimal medium SD, where *TSA1* deletion had a negligible impact on growth. When acetic acid is applied, again *TSA*1 deletion negatively affected growth in the T73 background but not in L2056 and EC1118 strains. Interestingly, *TRR1* deletion exhibited a stronger negative impact in the L2056 background compared to the other strains. A different stress was applied to rule out a general stress sensitivity. High ethanol (10%), a stress relevant during winemaking, was tested. Under these conditions, *TSA1* deletion increased tolerance to ethanol, with a mild effect observed in L2056 and slightly greater effect in T73, while no impact was detected in EC1118. In contrast, *TRR1* deletion mutant displayed slightly increased ethanol sensitivity in T73, but not in the others. These findings highlight the influence of genetic background on the phenotypic outcomes of deleting components of the peroxiredoxin-thioredoxin system under adverse conditions. Since T73 exhibited the most pronounced responses, it was selected for subsequent experiments.

### Tsa1 plays a role in acetic acid production and consumption during growth

To assess the impact of *TSA1* deletion on acetic acid metabolism during growth, T73 and T73 *tsa1*Δ strains were grown in Glucose Minimal Medium (GMM), enabling the monitoring of acetic acid dynamics across metabolic phases under laboratory standard conditions. Both strains exhibited comparable glucose uptake (Figure 3A) and ethanol production (Figure 3B) rates. While the absolute values for glucose consumption and ethanol production suggest a minor influence of Tsa1 on fermentative metabolism under these conditions, the overall yield remains unaffected. This is likely due to the slight growth defect observed in the *tsa1*Δ mutant strain (Supplementary Table 2). This indicates that Tsa1 has a slight influence on central carbon metabolism under these particular conditions.

**Figure 3.**
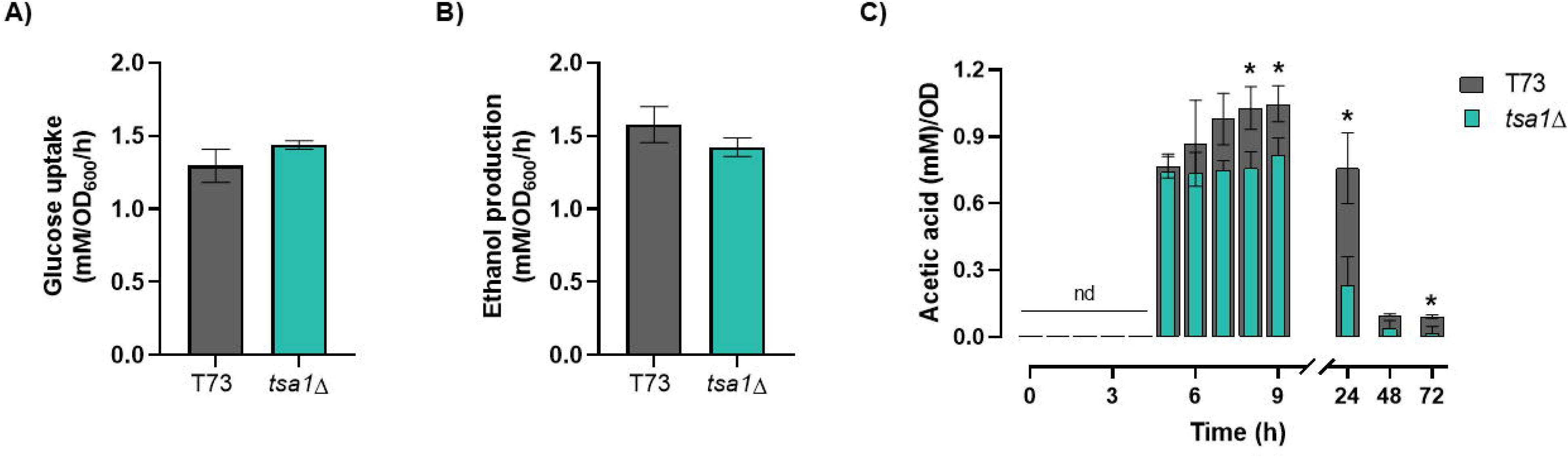
Tsa1 impacts acetic acid production and consumption in laboratory growth media in a wine yeast strain. A) Glucose uptake B) ethanol production and C) Acetic acid yields in GMM medium by T73 and T73 *tsa1*Δ strains. Experiments were carried out in triplicate, and the average and standard deviation are provided. Significant differences (*p < 0.05, Student’s t-test) between the mutants and their parental strains are shown. nd, not detected.

In contrast, acetic acid metabolism showed higher differences (Figure 3C). In the T73 strain, acetic acid was produced during growth and slightly increased to reach a maximum at the diauxic point of this strain. After that, its concentration decreased to very low levels when ethanol is fully consumed. Therefore, respiratory metabolism also uses acetic acid as a nutrient. The mutant strain exhibited a similar overall profile, but absolute acetate levels were lower after 8 hours of growth, the maximum concentration was also reduced, and its decline was faster, particularly at 24 hours. At the final point of the experiment (72h) these differences also remained significant. Therefore, Tsa1 seems to be involved in both acetic acid production and consumption in wine yeasts under laboratory conditions. In order to see whether this effect is conserved in a laboratory strain, the same experiment was carried out in the W303 genetic background (Supplementary Figure 1). The glucose uptake and ethanol production rates were similar, with no differences between wild type and *tsa1*Δ strains. The acetic acid yield profile in the *tsa1*Δ mutant was nearly identical to its parental strain, with no significant differences. These findings underscore the phenotypic differences between industrial and laboratory strains, emphasizing the importance of validating laboratory-derived claims in industrial wine strains. Based on these observations under laboratory conditions, we next aimed to investigate whether Tsa1 really does not influence the acetate-driven extracellular pH homeostasis in wine strains, as previously reported (Palkova et al., 1997).

### Media acidification is impacted by peroxiredoxin mutation

It has been described that during growth in solid media, yeast colonies acidify their surroundings through acetic acid excretion (Palkova et al., 1997). This phenomenon can be followed using a pH-sensitive dye, bromocresol purple, incorporated into the growth medium (Figure 4A). Drops of cultures from T73, *tsa1*Δ and *trr1*Δ strains were placed on YPD plates buffered at pH 6.5, and the acidification around the “giant colonies” was followed over two weeks. In the wild-type strain, the yellow halos indicating acidification started around day 8, persisted until day 10, and then the medium was re-alkalized by day 13. In the *tsa1*Δ mutant, the acid burst was delayed to day 10 but similarly re-alkalized by day 13.

**Figure 4.**
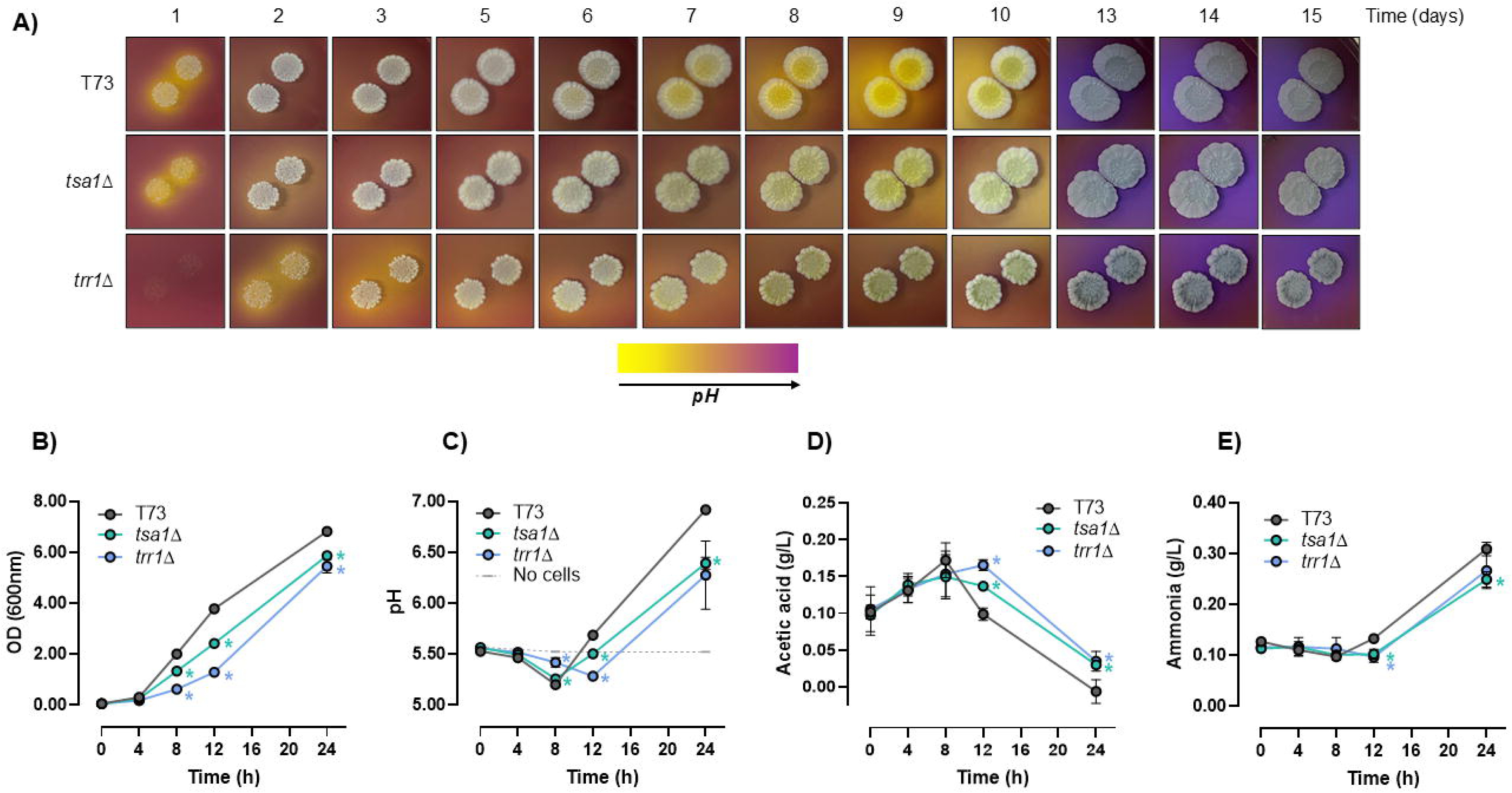
Deletion of *TSA1* reduces extracellular pH by maintaining higher acetic acid levels. The impact of *TSA1* deletion on external pH was measured with T73, T73 *tsa1*Δ and T73 *trr1*Δ strains in two media, YPD plates buffered at pH 6.5 with 0.01% bromocresol purple, BCP (A) and a liquid medium with 0.2% glucose as carbon source, pH 5.5 (B-E). A) Drops of both strains were placed on plates with BCP to obtain “giant colonies” and changes in external pH were monitored for 15 days. B) Growth measured as optical density (OD) at 600 nm. C) Extracellular pH. D) Acetic acid concentration. E) Ammonia concentration. Experiments in liquid medium (B-E) were carried out in triplicate, and the average and standard deviation are provided. Significant differences (*p < 0.05, Student’s t-test) between the mutants and their parental strains are shown.

In order to make a more quantitative analysis of pH variation, an alternative liquid culture approach was used (Figures 4B and C) in a medium with very low glucose (0.2%). Under these conditions, the shift from fermentation to respiration can be followed before 24 hours. Growth curves (Figure 4B) revealed that the T73 wild-type strain exhibited the fastest growth rate, followed by the *tsa1*Δ mutant, while the *trr1*Δ strain showed the slowest growth. These findings indicate that under low-glucose conditions the deletion of cytosolic peroxiredoxin-thioredoxin reductase system components has a significantly greater impact than in glucose-rich environments (Figure 2A). Both T73 and *tsa1*Δ had a fast drop in pH during the fermentative phase, increasing later when cells are under respiration (Figure 4C). However, the pH increased in the wild-type strain more efficiently than in the *tsa1*Δ mutant, which maintained lower pH levels at longer times. The *TRR1* deletion strain exhibited delayed medium acidification and a slower subsequent rise in pH, reaching a level similar to the *tsa1*Δ mutant. Acetic acid was measured along the growth curve (Figure 4D). All strains showed an increase in acetic acid up to the 8 hours that explain the drop in pH during fermentation. After that, acetic acid declined rapidly in the wild type, but not so in the mutants, indicating that the thioredoxin-peroxiredoxin system impacts acetic acid assimilation in post-diauxic conditions.

On solid plates, the re-alkalization of the media has been linked to ammonia production (Palkova et al., 1997). When ammonia was measured in liquid cultures (Figure 4E), a later burst in ammonia production could contribute to the pH rise, but this event was not linked to either Trr1 or Tsa1 activity.

### Genetic interaction between Tsa1 and aldehyde dehydrogenases

To understand how Tsa1 regulates acetate metabolism in wine yeasts, we explored its potential influence on the primary aldehyde dehydrogenases. These enzymes are at the core of acetic acid production, with mitochondrial Ald4 primarily contributing during respiratory metabolism, while cytosolic Ald6 plays a dominant role under fermentative conditions. *ALD4* and *ALD6* deletion mutants were combined with *TSA1* knock-out. To facilitate the construction of double mutants, the haploid wine strain C9 (Walker et al., 2003) was used in this section. First, the growth and acetate production of single and double mutants were tested in YPD plates containing the pH-sensitive dye BCP (Figure 5A). In this genetic background, the acid burst was more prolonged in time. *tsa1*Δ and *trr1*Δ mutants showed a lower level of acid production. As expected, the acidification was seriously impaired in the *ald4*Δ strain, confirming that Ald4 is the main acetic acid production under these post-diauxic conditions. The deletion of *ALD6* also affected acid production, albeit to a lesser extent. The double mutant *tsa1*Δ*ald4*Δ strain displayed an acidification pattern similar to that of the *ald4*Δ single mutant but with a slight delay in media re-alkalization. Interestingly, *tsa1*Δ*ald6*Δ showed evidence of an additive effect at longer times.

**Figure 5.**
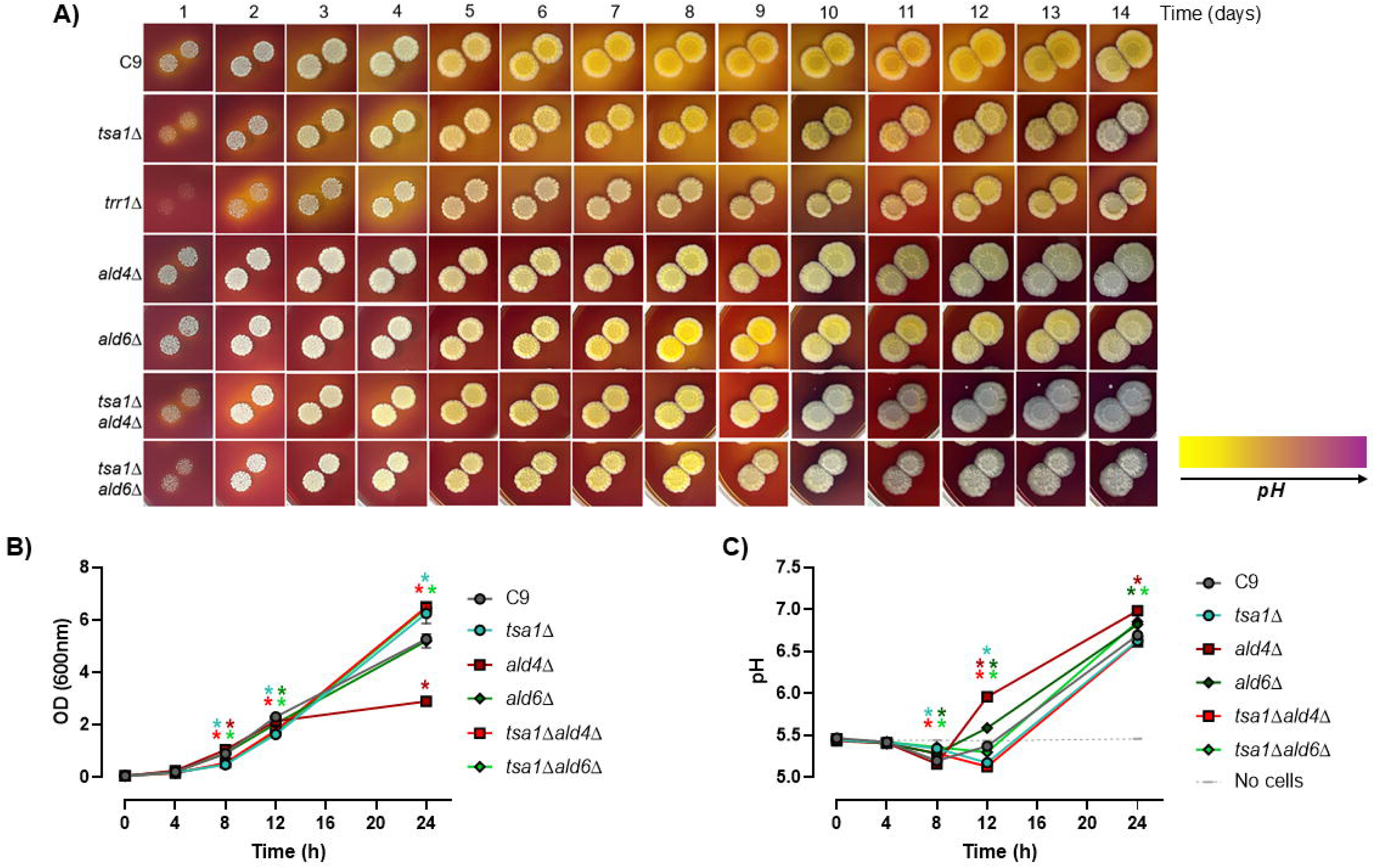
*TSA1* interacts genetically with *ALDH* in terms of medium acidification. Similar experiments of Figure 4 were performed with C9, *tsa1*Δ, *trr1*Δ, *ald4*Δ, *ald6*Δ, *tsa1*Δ*ald4*Δ and *tsa1*Δ*ald6*Δ strains. A) Color evolution in YPD with BCP for 14 days. B) Growth measured by OD_600_ in liquid medium with glucose 0.2%. C) pH in the conditions described in B). Experiments in liquid medium (B-C) were carried out in triplicate, and the average and standard deviation are provided. Significant differences (*p < 0.05, Student’s t-test) between the mutants and their parental strains are shown.

Growth in low carbon can provide a clearer perspective of this interaction (Figures 5B and 5C). Under these conditions, the growth profile of the *ald6*Δ strain was very similar to the parental strain (Figure 5B). In contrast, the *ald4*Δ strain grew similarly to the wild type during fermentative metabolism but reached lower densities at later stages, when respiration is required. Therefore, Ald4 is relevant when mitochondrial metabolism is fully active. The *tsa1*Δ mutant exhibited slower growth than the wild type during the initial phase but reached a higher optical density at the end of the experiment. Interestingly, double mutants involving *tsa1*Δ displayed phenotypes that aligned closely with the *tsa1*Δ single mutant. Moreover, the growth defect of the *ald4*Δ mutant was completely suppressed by *TSA1* deletion, suggesting a compensatory interaction. The pH dynamics (Figure 5C) further clarified these relationships. As expected, the *ald4*Δ mutant exhibited the most pronounced impact, with a significant pH increase at later time points corresponding to respiratory growth. During early fermentative metabolism, acidification in the *ald4*Δ strain was unaffected. The *ald6*Δ mutant also influenced pH increase but to a lesser stent. The *tsa1*Δ mutant displayed a delay in acidification during early growth but eventually reached nearly normal pH levels. Remarkably, the *TSA1* deletion fully suppressed the pH phenotype of *ald4*Δ mutant. In the *tsa1*Δ *ald6*Δ double mutant, the acidification delay observed in the *tsa1*Δ strain persisted, but the final levels were comparable to those of the *ald6*Δ mutant. This indicates a partial suppression effect, dependent on the metabolic status of the cell. After characterizing the genetic interactions between *TSA1* and *ALD4/6* in standard laboratory medium, we extended our investigation to winemaking conditions. This fermentative and industrially relevant environment, where *S. cerevisiae* wine strains are typically employed, allowed us to assess the role of *TSA1* in regulating these enzymes under conditions that closely mimic actual wine production.

### Ald6 activity is controlled by Tsa1 during winemaking

Peroxiredoxin Tsa1 exhibits a multifaceted role in acetic acid production in a variety of environments. However, the main biotechnological impact of acetic acid in winemaking lies in its contribution to volatile acidity during grape juice fermentation, so a closer look was taken at this process. In order to perform molecular analysis of vinification, a standard synthetic grape juice (MS300) was used (Figures 6 and 7) for microvinifications with T73 and T73 *tsa1*Δ strains. Fermentation was followed by reducing sugar consumption (Figure 6A). Fermentation speed was higher in the parental strain, although the *tsa1*Δ mutant ultimately completed fermentation to dryness. Acetic acid production was measured during the 7 days that lasted T73 fermentation (Figure 6B). In the wild-type strain, acetic acid was rapidly produced from day 1 and remained high during fermentation. In contrast, the *tsa1*Δ mutant displayed reduced acetic acid production on day 1, with levels recovered at longer times of fermentation. The current knowledge indicates that the major cytosolic aldehyde dehydrogenase Ald6 is the main contributor to acetic acid production during wine fermentation. Ald6 activity, measured in the presence of magnesium and NADP^+^ as a cofactor, supported this conclusion (Figure 6C). In the parental T73, Ald6 activity peaked in the first day, corresponding to the rapid acetic acid production observed at this time. This activity subsequently decreased and stabilized at lower levels during the rest of the fermentation. *TSA1* deletion caused a decrease in activity on day 1, which matches with the initial lower production of acetic acid (Figure 6B) and points to Tsa1 as a regulator of Ald6 activity. However, this activity in the mutant approached wild-type levels later on. To evaluate mitochondrial ALDH, its activity was measured in the presence of K^+^ ions and NAD^+^ as a cofactor (Figure 6D). As expected under fermentative conditions, Ald4 activity was substantially lower than Ald6. In the T73 strain, it was negligible on day 1 when most acetic acid is produced but increased at longer times, albeit remaining significantly lower than the cytosolic activity. *TSA1* knockout did not notably impact this activity, as both wild-type and mutant strains showed similar patterns.

**Figure 6.**
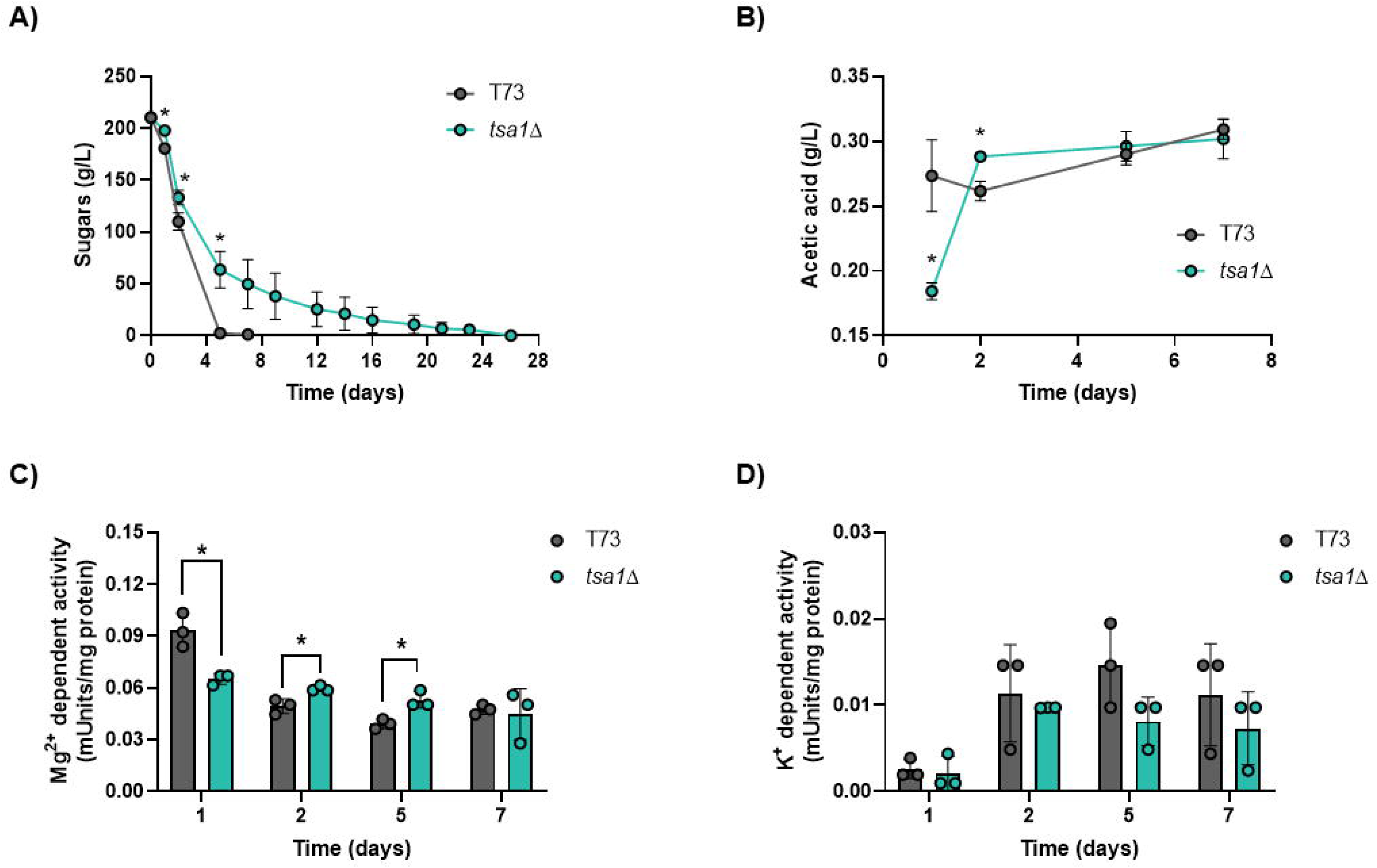
Tsa1 controls Ald6 activity during winemaking. Fermentations on synthetic grape juice MS300 by T73 and T73 *tsa1*Δ strains were performed. A) Fermentation evolution was measured as reducing sugar consumption. B) Acetic acid production during the first 7 days of fermentation. C) Cytosolic ALDH activity Ald6 during the time points described in panel B). D) Mitochondrial ALDH activity Ald4 during the time points described in panel B). Experiments were carried out in triplicate, and the average and standard deviation are provided. Significant differences (*p < 0.05, Student’s t-test) between the mutant and the wild-type strains are shown.

**Figure 7.**
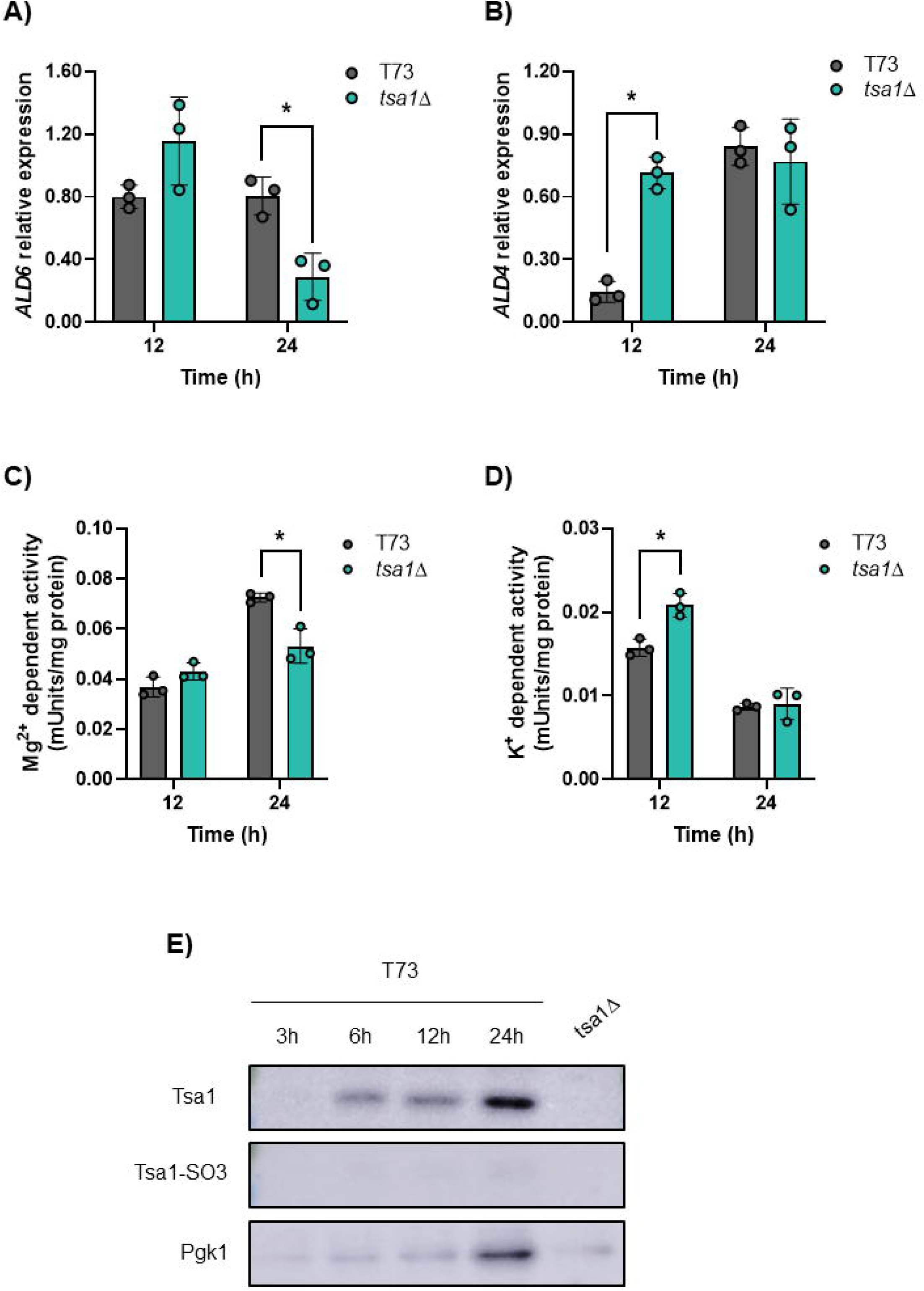
Tsa1 regulates early *ALD4/6* expression and activity. Fermentations on synthetic grape juice MS300 by T73 and T73 *tsa1*Δ strains were performed, and gene expression was measured in the first 24h. A) *ALD6* and B) *ALD4* mRNA levels measured by qPCR. C) Mg^2+^ dependent cytosolic ALDH activity at the same time points. D) K^+^ dependent cytosolic ALDH activity at the same time points. E) Western blot showing total and sulfenylated levels of Tsa1 in wild-type T73 strain. A lane containing the protein extract from the T73 *tsa1*Δ strain at 24h of fermentation was included as a negative control. Pgk1 levels were used as a loading control. Experiments were carried out in triplicate, and the average and standard deviation are provided. Significant differences (*p < 0.05, Student’s t-test) between the mutant and the wild-type strains are shown.

### Tsa1 impacts gene expression

As the enzymatic activity of ALDH is impacted by *TSA1* deletion, a closer look was taken to its potential mechanism. To do so, mRNA levels of the main aldehyde dehydrogenase genes *ALD6* and *ALD4* and the corresponding enzymatic activities were analyzed at the beginning of the vinification, where the main differences were recorded (Figure 6). Quantitative PCR (qPCR) revealed distinct patterns of *ALD6* and *ALD4* expression over time (Figures 7A and 7B). For *ALD6*, mRNA levels were comparable between the wild-type and *tsa1*Δ strains at 12 hours but were significantly reduced in the mutant at 24 hours (Figure 7A). This is in agreement with Mg^2+^-dependent cytosolic ALDH enzymatic activity (Figure 7C). *ALD4* gene expression exhibited a more complex profile. *TSA1* deletion increased *ALD4* levels at 12 hours but this difference was lost at 24 hours (Figure 7B). Mitochondrial ALDH enzymatic activity mirrored this trend with elevated activity at 12 hours in the mutant but no significant differences at 24 hours between strains (Figure 7D). Under this environment *ALD4* is affected by glucose repression, which may mask any longer-term effects of *TSA1* deletion on mitochondrial ALDH activity. Due to this reason, the absolute levels of the mitochondrial activity is lower than the cytosolic one, causing only a small contribution to the final levels of acetate.

Finally, the levels of Tsa1 protein itself during the first day of wine fermentation were measured by Western blot (Figure 7E). Tsa1 was barely detectable at 3 hours after inoculation but increased significantly during the next hours to reach a maximum at 24h. The *tsa1*Δ strain was used as a control of the specificity of the antibody. Oxidative stress was assessed by detecting hyperoxidized (sulfinylated) Tsa1 using a specific antibody. Tsa1 sulfenylated form indicates a strong oxidative stress where the peroxiredoxin has been damaged and cannot be recycled by the redox-controlling machinery. No hyperoxidized Tsa1 was observed, suggesting the absence of oxidative damage during the initial stages of fermentation. This indicates that the regulatory role of Tsa1 in acetic acid metabolism and ALDH activity is not primarily driven by its antioxidant function.

## Discussion

The role of peroxiredoxin Tsa1 on acetic acid metabolism was described in detail in this work. The findings reveal a complex relationship, as *TSA1* deletion impacts acetic acid production and consumption in diverse and sometimes apparently contradictory ways, depending on the growth media and conditions. These conditions include both laboratory and industrially relevant settings, where exogenous pH is also significantly impacted.

Acetic acid is a critical compound in winemaking, as it constitutes the main chemical species behind the undesirable volatile acidity. In a prior work from our laboratory focused on the role of Tsa1 during biomass propagation, preliminary experiments during grape juice fermentation showed a decrease in acetic acid in the deletion mutant (Garrigós et al., 2020). Despite being a key issue in winemaking, acetic acid production and management during fermentation is not completely elucidated (Vilela-Moura et al., 2011). Factors such as grape juice composition, particularly nitrogen and vitamin content, initial sugar concentration, as well as physical variables like temperature and aeration, all play significant roles on volatile acidity. To add more complexity to this issue, it is well known that the genetic background strongly influences acetic acid production (Patel and Shibamoto, 2002; Shimazu and Watanabe, 1981; Torrens et al., 2008), so this is a factor that is taking into account when new strains are isolated from the wild, and when the commercially available strains are marketed. We have observed variability on the impact of the deletion of peroxiredoxin *TSA1* and thioredoxin reductase *TRR1* in the growth on acetate as a carbon source and tolerance to high levels of acetic acid (Figure 2). These results suggest that redox controlling machinery may be contributing to that genetic heterogeneity observed in acetic acid metabolism and tolerance, an aspect that may warrant consideration in future studies and industrial applications. For experimental consistency, much of this work has focused on the commercial T73 strain. However, given the vast genetic variability of wine *S. cerevisiae* strains, conclusions drawn here may not apply universally across all strains (Borneman et al., 2016).

This study highlights that the *tsa1*Δ mutant impacts both the production and the consumption of acetic acid. Even in the catabolic side of metabolism, the effect of *TSA1* deletion is highly dependent on the initial metabolic conditions. For instance, the *tsa1*Δ deletion mutant shows accelerated acetic acid consumption during the post-diauxic phase of a 1% glucose culture (Figure 3) but exhibits elevated acetic levels when the culture is mainly respiratory, with just 0.2% glucose (Figure 4D). This underscores the role of the thioredoxin-peroxiredoxin system in sensing and adapting cellular metabolic status to regulate metabolism. While this study primarily focuses on enzymes responsible for acetic acid production, the catabolic side of metabolism is also crucial. When acetate is the sole carbon and energy source, it is metabolized to acetyl-CoA by acetyl synthase proteins. As *tsa1*Δ mutants display impaired growth on acetate (Figure 2A), and since Tsa1 is a cytosolic protein, it may exert its regulatory influence over the Acs1 isoform, which mainly provides cytosolic acetyl-CoA for lipids biosynthesis. We have previously observed an impact of thioredoxin deletion in the lipid biosynthesis that may point in the same direction (Picazo et al., 2019). A deficiency in lipids synthesis could partially explain the poor growth of the mutant in these conditions. Furthermore, acetyl-CoA generated by these enzymes is required not only for lipid synthesis but also for chromatin silencing. This dual role could suggest that alterations in acetyl-CoA metabolism may indirectly affect gene expression, potentially contributing to the range of phenotypes observed in the *tsa1*Δ mutant.

Production of acetic acid in fermentative conditions primarily arises from the oxidation of part of the acetaldehyde produced as an intermediary in alcoholic fermentation by a variety of aldehyde dehydrogenases. Under winemaking conditions, the main cytosolic ALDH (Ald6) is the predominant contributor to acetic acid production, while the mitochondrial isoform (Ald4) is less active (Remize et al., 2000; Saint-Prix et al., 2004). This was confirmed in our experiments (Figure 6), where the reduction in acetic acid in the *tsa1*Δ mutant correlated with a decrease in the magnesium and NADP^+^-dependent ALDH activity that links to the cytosolic Ald6 isoenzyme. The genetic interaction analysis revealed a more intricate regulatory picture (Figure 5). Interestingly, the absence of *TSA1* suppressed the phenotype of both *ALD4* and *ALD6* deletion mutants. Individually, these mutants increased the pH due to lower acetic acid production in respiratory laboratory conditions. The observed alkalinization linked to ammonia secretion has been associated with cellular communication between cells inside a colony (Palkova et al., 1997). However, our data shows that Tsa1 had no direct effect of ammonia production. Instead, its impact was linked to the acid burst associated with acetic acid. This phenomenon, previously connected to mitochondrial superoxide dismutase (Sod2) through Ald4 (Baron et al., 2013), has now been shown to also involve peroxiredoxins under our respiratory conditions, placing Tsa1 as a key regulatory node. Our previous work shows that cytosolic thioredoxin mutant regulates cytosolic Sod1 too (Picazo et al., 2023). Superoxide dismutases (SODs) transform superoxide into hydrogen peroxide that can act as a second messenger in different processes (Reddi and Culotta, 2013). Tsa1 can be the sensor of some signals generated by SODs to control both aldehyde dehydrogenases.

Peroxiredoxins, particularly Tsa1, are increasingly recognized as regulators of cellular processes rather than merely detoxifying enzymes. Evidence from this study supports this perspective, as the absence of hyperoxidation in Tsa1 catalytic cysteine during winemaking (Figure 7) indicates its regulatory role is triggered by more subtle inputs than a stressful rise in reactive oxygen species (ROS). The mechanism of action of Tsa1 can be direct or indirect. It has been known that peroxiredoxins modulate the activity of enzymes and transcription factors by regenerating oxidized cysteine residues. For instance, the oxidative-responsive transcription factor Yap1 is activated via the formation of a disulfide bond, which increases its nuclear localization. This redox regulation has been linked to the ability of peroxiredoxin to relay their oxidative status (Tachibana et al., 2009). Mutations in *YAP1* have previously been associated with altered acetic acid production during winemaking (Cordente et al., 2013), suggesting that Tsa1 may regulate gene expression involved in acetic acid metabolism through a similar mechanism. Specifically, Tsa1 might directly regulate the redox status of cysteines residues of one or several enzymes involved in acetic acid synthesis and/or degradation. Among them, enzymes sharing cytosolic localization with Tsa1 are the most likely candidates, being the aldehyde dehydrogenase Ald6 a prominent example. Global analysis suggest that Tsa1 may bind to Ald6 under specific conditions, and that Ald6 contains a cysteine residue susceptible to redox regulation (Seisenbacher et al., 2023). However, direct regulation alone cannot account for the modulation of mitochondrial ALDH Ald4, making an indirect mechanism that also encompasses transcriptional regulation the most reasonable explanation. A likely candidate for this mechanism is the ability of Tsa1 to modulate the nutrient signaling pathway through protein kinase A (PKA) (Roger et al., 2020). The cAMP-dependent PKA pathway promotes growth and protein synthesis under glucose-rich conditions. Tsa1 has been shown to stimulate sulfenylation of cysteine residues in the PKA catalytic subunit, thereby repressing it. PKA activity regulates numerous metabolic pathways and represses general stress response, acting at both transcriptional and post-transcriptional levels. This multifaceted regulation provides several potential ways for controlling acetic acid metabolism. For instance, in previous work, we described that *ALD4* was regulated by the general stress response transcription factors Msn2/4, direct targets of PKA (Aranda and Ml, 2003). The net impact of *TSA1* deletion on acetic acid levels is highly dependent on the growth and metabolic status of the cell, leading to seemingly contradictory outputs. One potential explanation for this erratic behavior may lay in the fact that the oxidation cycles of peroxiredoxins, including Tsa1, have been linked to metabolic oscillations (Causton et al., 2015). Hence, the absence of Tsa1 might exert opposite effects at different phases of these circadian rhythms, such as yeast respiratory oscillation. Therefore, there is still much work ahead to fully grasp the nuances with which the redox regulatory machinery influences metabolism. Addressing this challenge will not only improve our understanding of these processes but will also provide innovative strategies to modulate the production of relevant molecules during industrial yeast fermentations.

## Author contributions

VG, JCE, EM and AA contributed to the study conceptualization and design. VG elaborated all figures. VG, CP and LD performed the experiments and the formal analysis. JCE and AA supervised the project. VG and AA wrote the first draft of the manuscript. All the authors have read and edited the final manuscript. JCE, EM and AA contributed to funding acquisition and project management.

