## Supplementary Table 1 for "Cytosolic peroxiredoxin Tsa1 influences acetic acid metabolism and pH homeostasis in wine yeasts"

**Supplementary Table 1**. List of the strains used in this study.

| **Yeast Strain** | **Genotype / Plasmid added** | **Source** |
| --- | --- | --- |
| L2056 | Commercial Wine Strain Lalvin Rhône 2056™ | Lallemand Inc. |
| L2056 *tsa1*∆ | L2056 *tsa1*::*loxP* *tsa1*::*KanMX* | Garrigós et al., 2020 |
| L2056 *trr1*∆*/TRR1* | L2056 *trr1*::*KanMX* *TRR1* | Garrigós et al., 2020 |
| L2056 *tsa1*∆ | L2056 CRISPR/Cas9 *tsa1*∆ | This study |
| L2056 *trr1*∆ | L2056 CRISPR/Cas9 *trr1*∆ | This study |
| T73 | Commercial Wine Strain Lalvin T73™ | Lallemand Inc. |
| T73 *tsa1*∆ | T73 CRISPR/Cas9 *tsa1*∆ | This study |
| T73 *trr1*∆ | T73 CRISPR/Cas9 *trr1*∆ | This study |
| EC1118 | Commercial Wine Strain Lalvin EC1118™ | Lallemand Inc. |
| EC1118 *tsa1*∆ | EC1118 CRISPR/Cas9 *tsa1*∆ | This study |
| EC1118 *trr1*∆ | EC1118 CRISPR/Cas9 *trr1*∆ | This study |
| W303 | MAT a, *ADE*, *LEU*, *HIS*, *TRP*, *URA* | Jennifer C. Ewald laboratory |
| W303 *tsa1*∆ | W303 *tsa1*::*loxP* | This study |
| C9 | C9 Mat a *ho*::loxP | Walker et al., 2003 |
| C9 *tsa1*∆ | C9 *tsa1*::*KanMX* | Picazo et al., 2018 |
| C9 *ald4*∆ | C9 *ald4*::*KanMX* | Own yeast collection |
| C9 *ald6*∆ | C9 *ald6*::*KanMX* | Own yeast collection |
| C9 *tsa1*∆ *ald4*∆ | C9 *tsa1*::*loxP ald4*::*KanMX* | This study |
| C9 *tsa1*∆ *ald6*∆ | C9 *tsa1*::*loxP ald6*::*KanMX* | This study |
