## Supplementary figures and images for "Cytosolic peroxiredoxin Tsa1 influences acetic acid metabolism and pH homeostasis in wine yeasts"

### Supplementary Figure 1

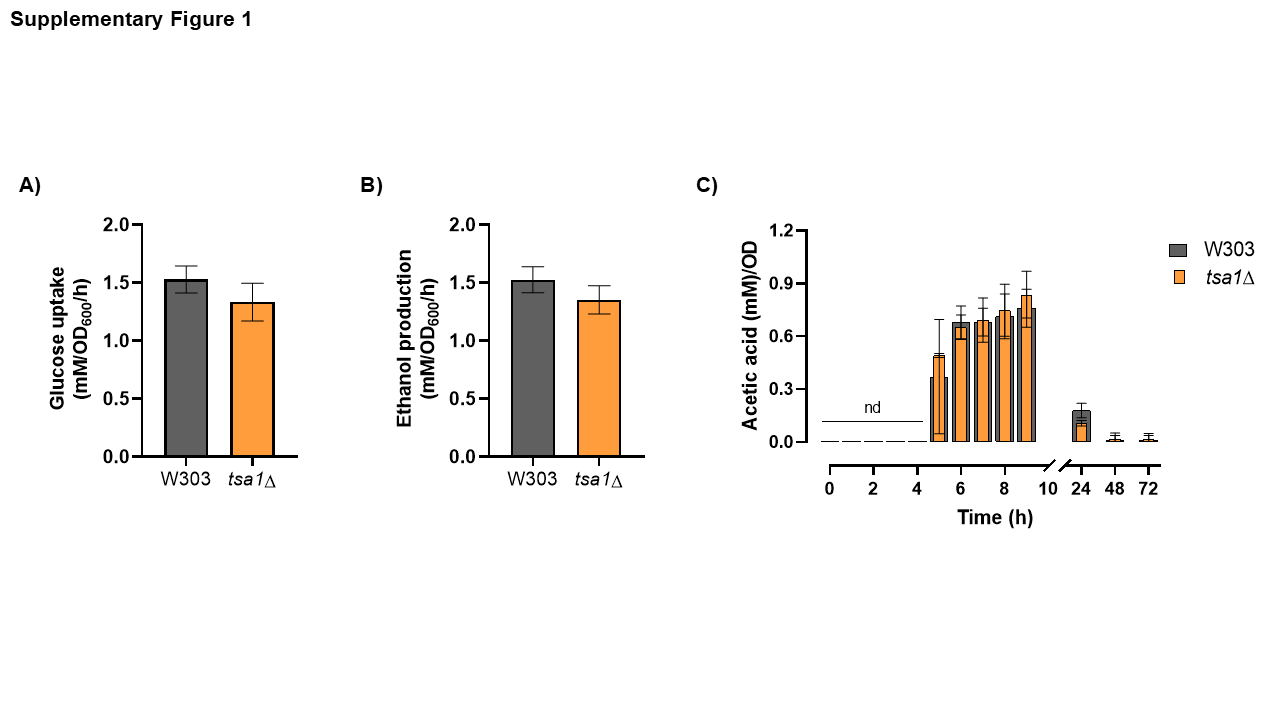
